# Validation of shoe sole dust as a microbial sampler reveals distinct fungal and bacterial responses to nearby vegetation

**DOI:** 10.64898/2026.03.17.712241

**Authors:** Sadia Marjia Ferdous, Pekka Taimisto, Emma Musakka, Taina Siponen, Martin Täubel, Bridget Hegarty

**Affiliations:** Department of Civil and Environmental Engineering, Case Western Reserve University, Cleveland, Ohio, USA; Department of Public Health, Finnish Institute for Health and Welfare, Kuopio, Finland; Institute of Public Health and Clinical Nutrition, University of Eastern Finland, Kuopio, Finland; Department of Civil Engineering, Aalto University, Espoo, Finland

**Keywords:** Microbial exposure, biodiversity, microbial sampling, amplicon sequencing, greenness

## Abstract

Urbanization-driven environmental change has significant implications for human health and well-being. However, studies have found differing patterns in microbial diversity along urbanization gradients; and it remains unknown whether this reflects methodological limitations or genuine ecological complexities. Resolving these inconsistencies requires innovative, reproducible methods that accurately reflect human contact with environmental microbiota. In this study, we have validated a new method for assessing environmental microbial exposure by measuring microbiota from particulate matter collected from shoe soles and studied the influence of vegetation at different proximities. Through repeated walks on routes along an urbanization gradient in Finland, we show that left and right shoe sole dust from the same walk and same route represent more similar microbial communities compared to different walks and routes. We found that bacterial biomass and diversity were best predicted by Normalized Difference Vegetation Index (NDVI, as a measure of greenness) immediately surrounding the walking path, whereas fungal communities responded to broader landscape-scale greenness (100m–1km), suggesting that bacteria and fungi are governed by different dispersal processes. Importantly, NDVI explained these differences in diversity more effectively than simple classifications of the path based on its substrate and whether it was in a rural or urban setting. Shoe sole dust sampling offers a simple, effective, and reliable approach for evaluating microbial exposures, capturing scale-dependent microbial responses to vegetation, and enabling more robust epidemiological studies on the health effects of greenness and environmental biodiversity.

## Introduction

By 2050, over two-thirds of the global population is projected to live in urban areas, dramatically reshaping natural landscapes and exposing billions to the health consequences of biodiversity loss and environmental degradation.^1^ While urbanization drives infrastructure development and economic growth, it also contributes to habitat loss, reduced biodiversity, and increased pollution, with negative consequences for human health and wellbeing. Nature exposures and biodiversity are increasingly connected with a broad range of health benefits, including improvements in cardiovascular health, mental wellbeing, physical activity, sleep quality, and metabolic outcomes such as type 2 diabetes^2–5^. According to the biodiversity hypothesis, early-life exposure to diverse environmental microbiota supports allergic health by shaping essential immunoregulatory pathways^6,7^. Therefore, the rising prevalence of chronic pro-inflammatory conditions, including asthma and allergies, may be linked to reduced biodiversity and limited nature contacts resulting from urbanization^8–11^. Complicating our understanding of the connections between urbanization and health, both protective and pre-disposing effects have been shown between *environmental greenness* (e.g., residential greenness and land use) and childhood allergic disease risk^12–18^. Suggesting that not all environmental greenness may lead to equivalent microbial exposures, while many studies have found that urbanization decreases microbial diversity^19–24^, there have also been observations of the opposite trend^25^.

Resolving whether the inconsistencies between studies reflects a true biological signal or methodological differences requires reproducible approaches that accurately reflect human contact with environmental microbiota during outdoor activities. In theory, air samples should be used to assess airborne microbial exposure. In practice, air sampling has multiple drawbacks for large cohort studies, including low biomass, short sampling windows, and expensive equipment. Indoor microbial exposures are typically collected with settled dust because these methods are simple for study participants to collect themselves and previous studies in residential settings have shown settled dust to be a reasonable surrogate^26–30^. Somewhat surprisingly, there is no established, comparable approach for assessing microbial exposure during outdoor activities. Most studies exploring associations between health outcomes and outdoor greenness, land use patterns, or biodiversity only consider the presence of the green space in defined buffers around the residential address, rarely considering the frequency of outdoor visits^31^. Studies linking outdoor nature visits to other health outcomes (e.g., cardiovascular or mental health) typically rely on questionnaires or diaries, though some have integrated GPS tracking with geospatial data to more accurately characterize environmental exposure during outdoor activities^32,33^. Population-level research that accurately assesses *biodiversity* in order to link exposure to immunological health is largely lacking^34^, in part because of the challenges of quantifying microbial exposures during outdoor activities^35^.

In this study, we address the need for more robust methods linking outdoor microbial exposures and health by introducing a novel approach: measuring microbiota from particulate matter collected from shoe soles. Previous studies have used this method for estimating bacterial exposures^36,37^, but not fungal. Here, we extend these works by assessing the reproducibility of these measures for the same walk and investigating how microbial communities vary with greenness. We have validated this approach through repeated walks on routes along an urbanization gradient. Our approach offers a promising framework for future population-based studies to help disentangle the contradictory findings thus far and reveal the mechanisms governing the connections between greenness, biodiversity, and human health.

## Materials and Methods

### 1. Sampling Campaigns

This study consisted of two sampling campaigns carried out in the Savo region in Eastern Finland. The first was a proof-of-concept study (“pilot campaign”) focused on a single 11 km route in the city of Kuopio. This route was walked twice per day for 17 days (34 walks) between March and May 2019. This route was conducted on paved walkways along major roads as part of another study to investigate human exposure to street dust during the spring ^38^. Shoe sole dust samples, as well as meteorological (temperature and relative humidity), air pollution (PM_2.5_ and PM_10_), and surface condition data (wetness, snow) were collected for each walk. The second study (“full campaign”) expanded to three different environments: urban-grey (a city area without larger green areas, with walks conducted exclusively on paved roads), urban-green (a city environment with large green areas, conducted on both paved roads and gravel paths), and rural-green (forest environment, conducted exclusively on paths with natural surface, such as forest soil, grass, and gravel). Two separate, non-overlapping 5 km routes were walked for each environment (Figure 1, Table 1, Figure S1). Each route was walked 5 times between July and September 2020, totaling 10 walks per environment, 5 walks per route, and 30 walks altogether. A trained researcher performed all the walks. We use the terms “environment”, “route”, and “walk” to represent the nested structure of our sampling.

**Figure 1:**
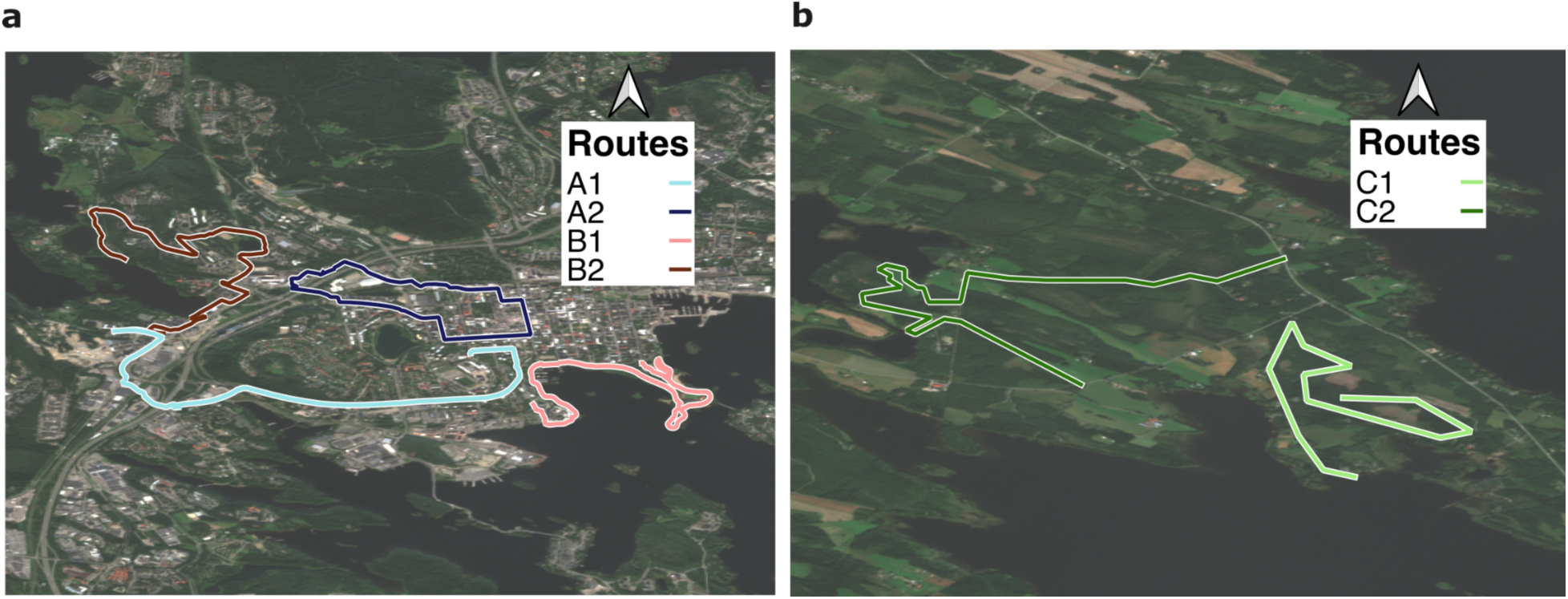
Spatial layout of the (a) rural and (b) urban sampling routes. Panel a includes both the urban-grey environment (A1-2 routes), and urban-green environment (B1-2 routes). Panel b contains the rural-green environment (C1-2 routes). Routes are color-coded as A1 (light blue), A2 (dark blue), B1 (light brown), B2 (dark brown), C1 (light green), and C2 (dark green).

**Table 1.** Description for walking surfaces and vegetation in close vicinity for different routes.

| Environment | Route | Length<br>(km) | Walking surface of the route<br>(% of total) |  |  |  | Vegetation type in close vicinity to the route<br>(% of total) |  |  |  |
| --- | --- | --- | --- | --- | --- | --- | --- | --- | --- | --- |
|  |  |  | Asphalt | Gravel | Forest<br>paths | Other | Street,<br>buildings,<br>minimal<br>green | Park-type<br>green<br>mixed<br>with<br>buildings | Dominantly<br>green areas<br>like parks | Forests |
| Urban-grey | A1 | 5.0 | 100 |  |  |  | 68 | 32 |  |  |
|  | A2 | 4.8 | 93 |  |  | 7 | 82 | 18 |  |  |
| Urban-green | B1 | 5.0 | 19.2 | 80.8 |  |  | 7 | 52.6 | 40.4 |  |
|  | B2 | 5.0 | 83.7 | 16.3 |  |  | 42.6 | 18.4 | 39 |  |
| Rural-green | C1 | 5.0 |  | 51.4 | 48.6 |  |  |  |  | 100 |
|  | C2 | 5.0 |  | 47.6 | 44.6 | 7.8 |  |  |  | 100 |

### 2. Shoe Dust Sampling

We used the same sampling process for shoe sole dust for both sampling campaigns. Shoes were new before the start of each sampling campaign. Before each walk the shoes were washed thoroughly with water, detergent and a brush for one minute. They were then sprayed with 70% ethanol and allowed to completely dry.

For each walk, samples were collected from the left and right shoes before and after the walk (4 samples total). The pre-walk swabbing samples were collected after cleaning, by swabbing the shoe sole for 15-20 seconds, including between the shoe profiles, with a sterile cotton swabbed wetted with a buffer (sterilized water plus 0.05% Tween 20). A separate swab was used for the left and right shoes. Swabbing was done in a clean facility. The cotton tip of the swab was cut into the DNA extraction tubes with sterile scissors and then deposited in a −20°C freezer. Shoes were transported to the start of the route in a clean plastic bag to avoid contamination before reaching the route. The walking route was monitored via GPS. At the end of the route, the shoes were put back in the plastic bag for transport to the lab. Post-walk swabbing was done in the same space and in the same way to pre-walk swabbing. Field blanks were also collected after each walk. We have provided a detailed protocol at protocols.io ^39^.

### 3. Air Sampling

In the full campaign, we also collected active air samples to investigate whether the microbial signal detected from shoe soles correlates with that of air samples collected during the walk. Active air sampling was conducted with Button Inhalable Aerosol samplers (SKC Inc. USA), and sampling lasted for the duration of the walk. One sample was collected for each route (six in total). Before sampling, Button samplers were pre-cleaned with 30 min sonication in water, followed by washing with Etax A, drying and assembly in a clean laboratory space. The samplers were transported in sterile plastic bags and handled wearing laboratory gloves to avoid contamination. Samplers were operated at a flow rate of 4 L/min (checked before and after walks with a Defender 510-H (Bios International Corporation, USA)), with a pump carried in a backpack and the inlet mounted at the height of the test subject’s breathing zone in the direction of walking. Button samplers were disassembled in a clean laboratory facility. Then, filters were stored in a Petri dish at −20°C until further processing. Button filters were placed in 2 mL Eppendorf tubes with 200 mg glass beads immediately before DNA extraction. For active air sampling, the field blank filters were collected by disassembling an unused Button Inhalable sampler.

### 4. DNA Extraction

DNA extraction was performed directly from the cotton swabs, or the Button filters. First, 400 µL lysis buffer of the Chemagic DNA Plant Kit (PerkinElmer Chemagen Technologie GmbH, Germany), and 10µl of salmon testis DNA (Sigma-Aldrich Co., USA) used as internal standard ^40^ to assess and correct for the presence of inhibitors and the performance of the DNA extraction, were added to the samples. This was followed by a bead-beating step for mechanical cell disruption ^41^ using a MiniBeadbeater-16 (Biospec Products, Inc., USA) at maximum speed for one minute. DNA was then purified on a KingFisher™ mL DNA extraction robot (Thermo Fisher Scientific, Inc., Finland), following the Chemagic DNA Plant Kit protocol. Reagent and negative controls as well as bacterial and fungal mock communities were included periodically in the DNA extraction process. DNA was stored at −20 °C until quantitative PCR and amplicon sequencing analyses.

### 5. Quantitative PCR (qPCR) analyses

Microbial concentrations in the samples were determined with qPCR on a QuantStudio™ 6 Flex Real-Time PCR System (Applied Biosystems, USA) utilizing previously published qPCR assays and PCR conditions ^42^ targeting Gram-positive and Gram-negative bacteria ^43^; as well as *Penicillium* spp., *Aspergillus* spp., and *Paecilomyces variotii* (PenAsp) ^44^, total fungal DNA (Unifung) ^45^; and internal standard salmon testis DNA ^40^. Due to inhibition issues affecting the Gram-positive side of the dual assay, only Gram-negative bacteria data are presented here.

### 6. Amplicon sequencing

Bacterial and fungal amplicon sequencing of samples from study 1 (“pilot campaign”) was done at a commercial sequencing service provider (LGC Genomics, Germany). DNA extracted from shoe sole debris as well as from blanks and control samples was shipped frozen to the service center. The V4 region of the bacterial 16S rRNA gene was amplified using 515F/806R primers ^46^ and ITS1 region of the Internal Transcribed Spacer (ITS) was amplified using ITS1F/ITS2 primers ^47^. Details on PCR conditions and the sequencing protocol have been described earlier (Dockx et al., 2021). Sequencing was then performed on an Illumina MiSeq with V3 chemistry resulting in paired-end reads with a length of 300 bp each. The libraries were demultiplexed using Illumina’s bcl2fastq v1.8.4 and all sequence reads processed with custom Python v2.7.6 scripts to sort them by sample, removing barcode and amplicon primer sequences. Adapter sequences were removed from the 3′ end of reads with a proprietary script discarding reads shorter than 100 bp.

Bacterial and fungal amplicon sequencing of samples from study 2 (“full campaign”) was performed at the CWRU Genomics Core. DNA extracted from shoe sole and air samples as well as from blanks and control samples was shipped frozen from Finland to Cleveland, USA. The V3/V4 region of the bacterial 16S rRNA gene was amplified using 341F/785R primers ^49^. The ITS2 region of the Internal Transcribed Spacer (ITS) was amplified using ITS3/ITS4 primers ^50^.

Sequencing was then performed on an Illumina MiSeq with V3 chemistry resulting in paired-end reads with a length of 250 bp each for both bacterial and fungal DNA. For both bacteria and fungi, samples with sufficient DNA were normalized to 10 ng/uL by the Genomics Core for library prep processing, following an optimized 16S Metagenomic Sequencing Library Preparation protocol with the 16S and ITS primers respectively. Details on the sequencing protocol has been provided previously (Leslie et al., 2025).

After sequencing, the reads from both the pilot and full campaigns were returned to the Hegarty Lab, then cleaned and processed using the DADA2 16S Pipeline Workflow v1.16 and DADA2 ITS Pipeline Workflow v1.8 ^52^. The DADA2 pipeline parameters were tuned to account for read length and required overlap to improve merging and error modelling of the sequencing reads. An ASV table with read counts was constructed after filtering, trimming and de-replication of reads. The sequences were annotated with the assignTaxonomy function (which uses a naive Bayesian classifier) in DADA2 based on the SILVA 138.2 ^53^ for bacteria and UNITE database ^54^ version 10 for fungi.

The sequencing for the pilot and the full study was done a few years apart, at different locations, and using different primer sets (Table S1.a-d). The main study extracted DNA samples that were stored longer and shipped from Finland to the USA for sequencing, which may affect the community composition ^55^. Other differences may result from samples being collected during different years and different seasons. For all these reasons and that the pilot route was not identical to any of the full-study routes, we did not compare the pilot samples directly to the full-study samples.

### 7. Greenness index

We downloaded the Copernicus Sentinel-2 Multispectral Instrument (MSI) Level-2A surface reflectance data at a 10-meter resolution for the study area for the date of August 5, September 3 and September 10, 2020 ^56^, corresponding to the timeframe of field sample collection. The images were processed in QGIS (v3.40.5) to calculate the Normalized Difference Vegetation Index (NDVI) using the red (Band 4) and near-infrared (Band 8) bands ^57^. NDVI values across all images were averaged to generate a single composite NDVI layer, which was then used for comparison with alpha diversity metrics. We calculated the NDVI for a buffer of 10m, 25m, 50m, 100m, 200m, 500m, 1km, 2km, 5km and 10km along the walking paths.

### 8. Data analysis and statistics

All analyses were done after rarefying the read counts to a depth of 6000 for bacteria and 5000 for fungi for the full campaign (Figure S2) and 8000 for both bacteria and fungi for pilot campaign to account for a high variability in sequencing depths across samples. After the DADA2 processing and rarefaction, we had 57 samples for the full campaign and 66 samples for the pilot campaign for both bacteria and fungi (after walk samples). Alpha diversity measures (observed richness and Shannon’s Diversity) were calculated using amplicon sequencing variants (ASVs) and the phyloseq (v1.48.0) package ^58^. Beta diversity of the microbial communities was quantified using the Bray–Curtis dissimilarity metric and visualized with non-metric multidimensional scaling (NMDS) using the ordinate function from the phyloseq package. ANCOM-BC2 ^59^ was used to test whether any bacterial or fungal taxa were differentially abundant between environments. The default settings were used, with the environment as a fixed effect. PERMANOVA using the adonis2 in the vegan package (v2.7.1) was used to evaluate the impact of the environmental parameters on the bacterial and fungal community. Distance-based Redundancy Analysis (db-RDA) ordination was used to visualize the bacterial and fungal community of the samples using the capscale function (vegan). Correlations between the environmental variables were performed using the cor function in the stats package (v4.4.0). Unless otherwise stated, the statistical tests were done using the wilcox.test and pairwise.wilcox.test from the stats package in R. As only two air samples (from the rural-green environment) were successfully sequenced, our ability to compare the air and walk samples was limited (Figure S3). The graphical abstract was created using Canva. All data analysis was done in R v4.4.0. Code and metadata files are available on the Hegarty Lab github page (https://github.com/HegartyLab/WITP). Sequences for both sampling campaigns are available from the NCBI Sequence Read Archive under the BioProject accession number PRJNA1380507.

## Results

### 1. Study overview

To validate using shoe sole dust as a microbial sampler, we conducted two complementary sampling campaigns. The first (“pilot campaign”) was focused on a single walking path in Kuopio, Finland; environmental data was collected for each walk. After validating the approach, we extended our sampling (“full campaign”) to 6 different paths in 3 types of environments: urban-grey (A), urban-green (B), and rural-green (C). Overall, the left and right shoe samples in each walk showed very similar bacterial and fungal communities that varied with environment (Figure 2). We quantify the similarity between the left and right samples and describe the changes in the microbial communities with route and environmental characteristics.

**Figure 2:**
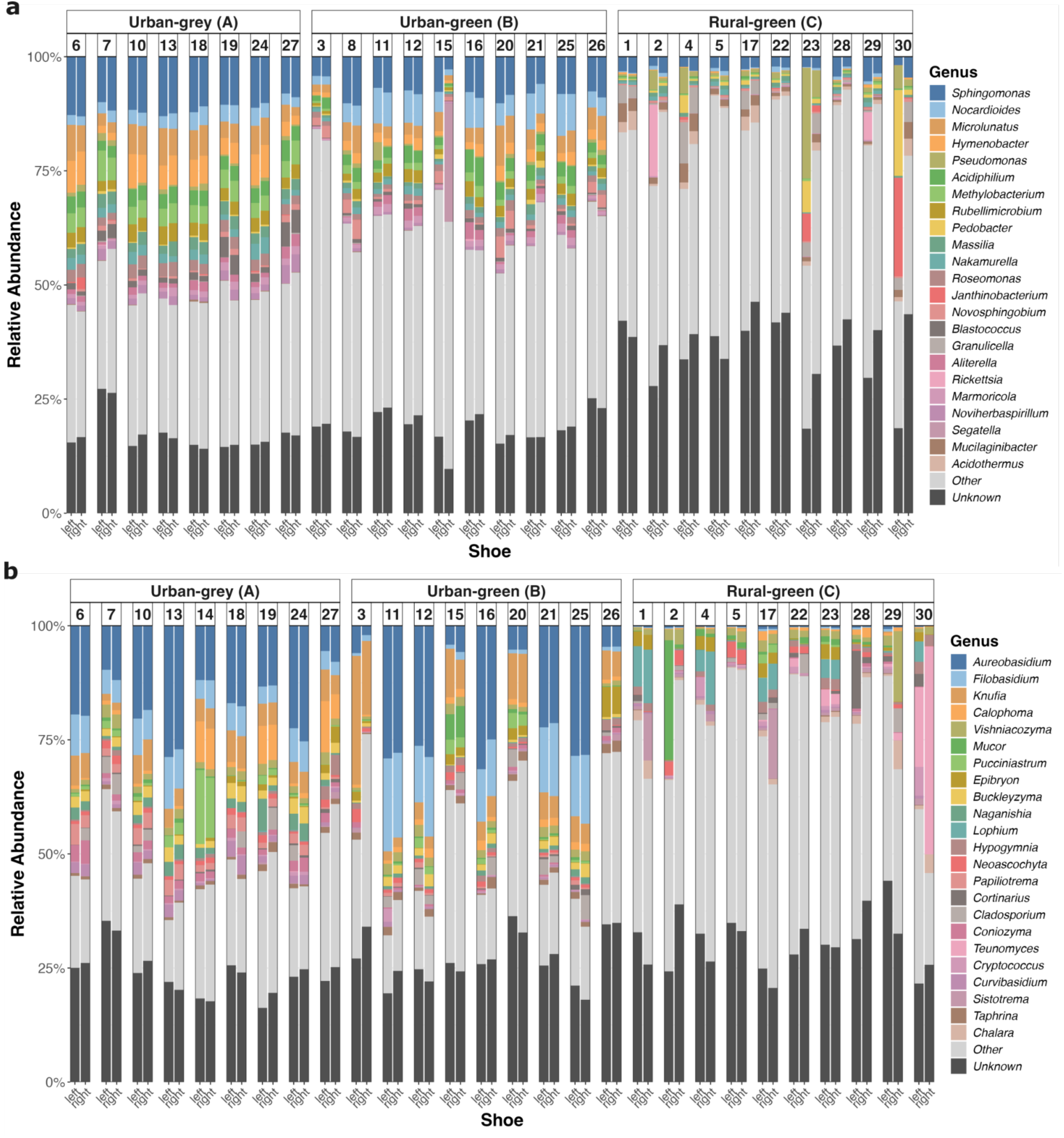
Genus level Top 20 taxa for (a) bacteria and (b) fungi by walk paired left vs right. Each bar is colored by genus. Samples are grouped by environment. The larger grey bars for the rural-green environment reflects the higher diversity of those routes. We excluded one sample here for not having the corresponding pair.

### 2. Left and right shoe sole dust capture microbial communities representative of the route

For both studies, paired left and right shoe samples were taken separately. Comparing the log_10_ transformed relative abundance of each genus between the left and right samples shows a clear linear relationship across all the environments, particularly for the most abundant taxa (Figure 3). This trend was stronger for the urban routes than the rural ones for fungi (R^2^ = 0.83 for urban-grey, 0.82 for urban-green, 0.73 for rural-green and p-value < 0.001 for all). The trend was similar for all the routes for bacteria (R^2^ = 0.86 for urban-grey, 0.85 for urban-green, 0.86 for rural-green and p-value < 0.001 for all).

**Figure 3:**
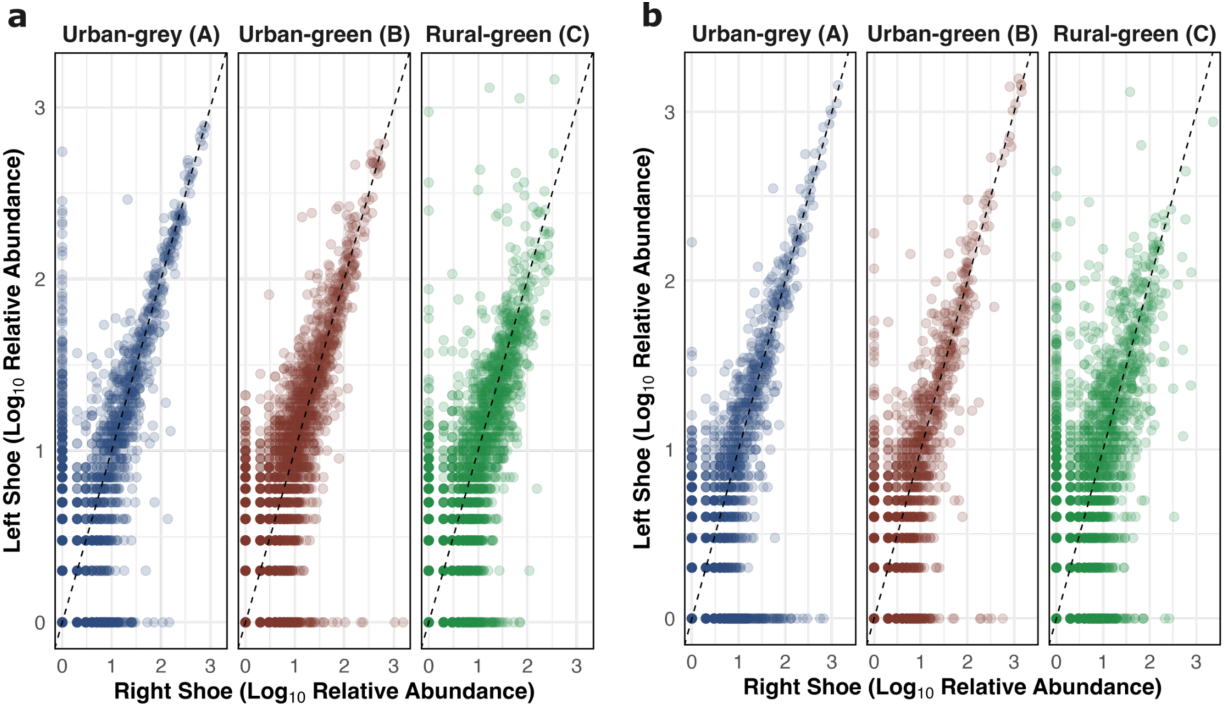
Scatterplot of the log_10_ transformed relative abundance of each genus for the left (x-axis) and right (y-axis) shoe across walks for (a) bacterial and (b) fungal communities. Relative abundances were log_10_ transformed with a pseudocount of 1 to accommodate zero values. Each point represents a genus in a given walk, and proximity to the 1:1 line indicates greater similarity in relative abundance between the left and right shoes.

For the pilot sampling campaigns, the left and right samples were significantly more similar when taken during the same walk as compared to a different walk (Figures 4 for genus level, Figures S4 for ASV level). Using Bray-Curtis Dissimilarities as a proxy for how similar two samples are, we found that left and right shoe bacterial and fungal communities are more similar during the same walk than during different walks (Figure 4; Wilcoxon test p-value < 0.001 for both). As the same route was walked twice daily over 17 days, we assessed whether the microbial community on the same path changed from the morning to afternoon. The microbial communities showed no statistically significant difference between the samples collected in the morning or afternoon of the same day (Table S2.a, S3.a), demonstrating that the difference between walks is due to differences between the sampling days.

**Figure 4.**
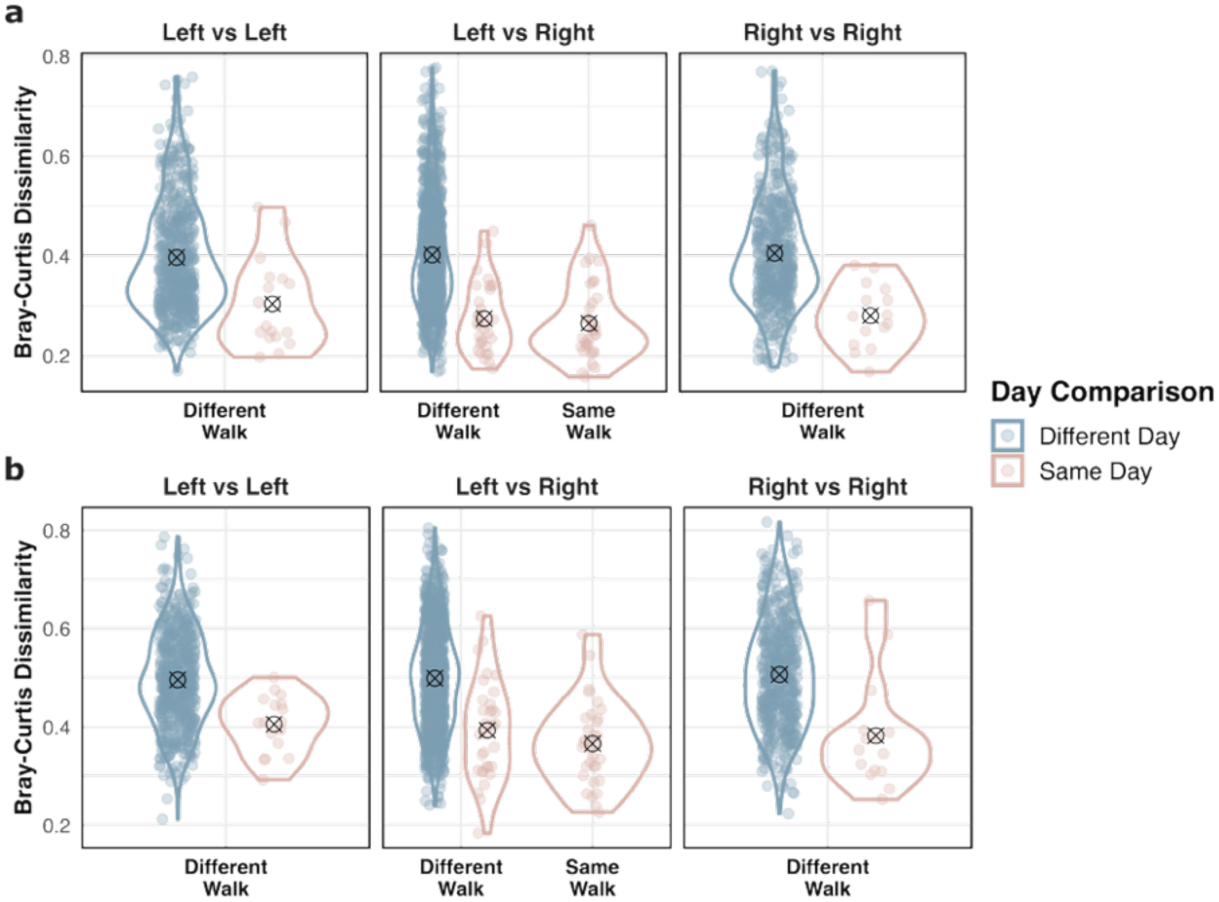
Bray-Curtis dissimilarities to compare the similarity between two samples’ microbial communities for the pilot campaign (genus level): (a) bacteria and (b) fungi. Violin plots are separated by walk and day. Points in pink are from the same day, representing the dissimilarity between paired left and right shoe samples from the same day (same walk) and those from the same day but not the same walk (different walk). Points in blue include all other dissimilarities (different day). The mean dissimilarity is represented by a circle with an x through it.

For the full-scale sampling campaign, we also compared the Bray-Curtis dissimilarities between samples taken from the same route (Figures 5 for genus level, Figure S5 for ASV level). We observed that the Bray-Curtis dissimilarities were much higher for samples taken from different route than for samples taken from the same route or the same walk (Wilcoxon test p-value < 0.001 for both). The dissimilarities for left vs right comparisons from different walks of the same route were similar to the dissimilarities for the left vs left and right vs right for each route. The total Gram-negative bacterial, fungal, and Pen/Asp gene copies for the full campaign were also more similar for the left and right shoe samples for the same walk and same route compared to the samples from different walks and routes (Figure S6-S7). Overall, the fungal communities on left and right shoe soles from the pilot data were less similar than the bacterial communities. However, this was not the case for the full-scale sampling campaign where both the fungal and bacterial communities showed similar ranges in their Bray-Curtis dissimilarities.

**Figure 5:**
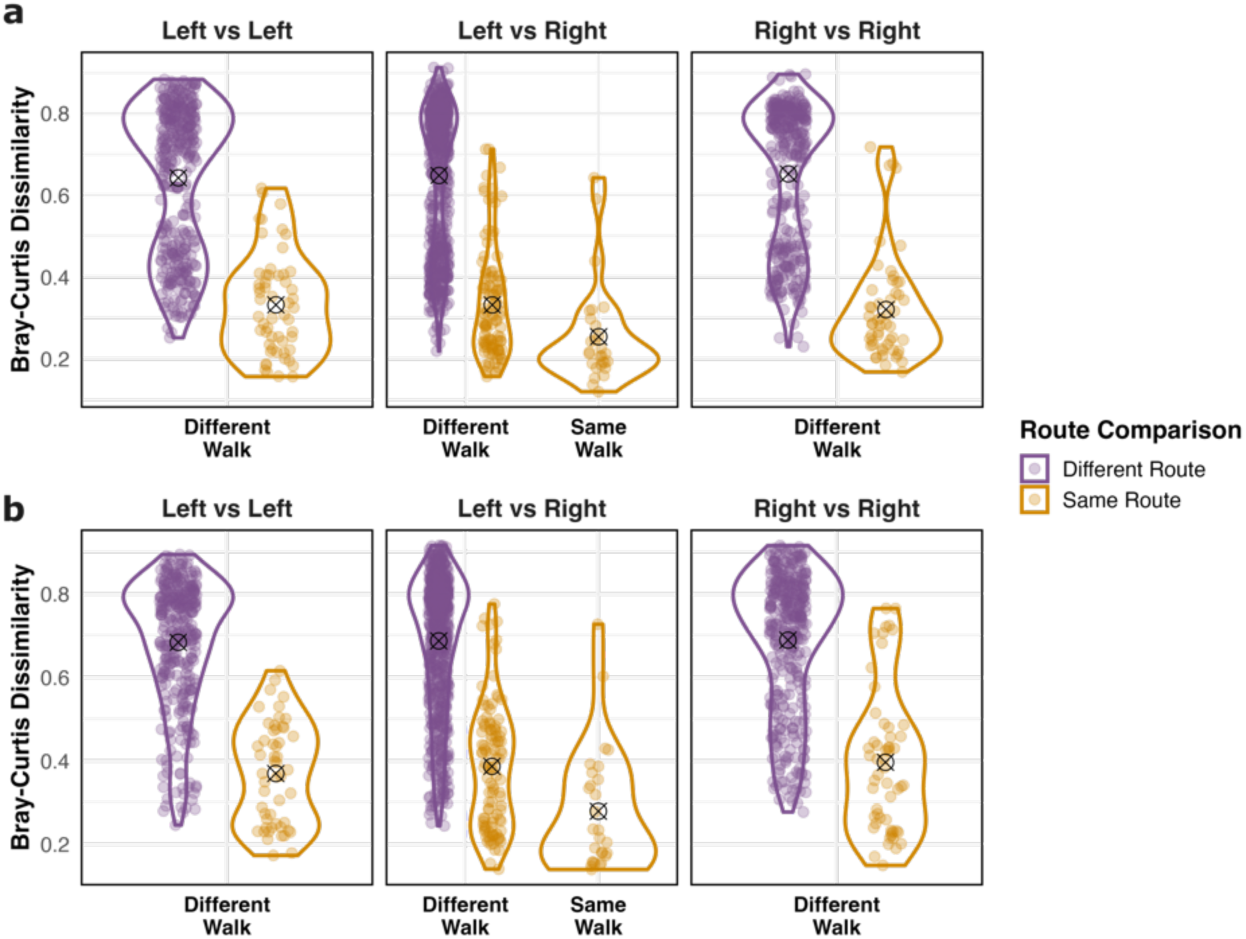
Bray-Curtis dissimilarities for the full campaign (genus level): (a) bacteria and (b) fungi. Violin plots are separated by walk and route. Points in gold are from the same route, representing the dissimilarity between paired left and right shoe samples from the same walk (same walk) and those from the same route but not the same walk (different walk). Points in purple include all other dissimilarities (different route). The mean dissimilarity is represented by a circle with an x through it.

### 3. Shoe sole dust microbiota varies with environmental conditions

In the pilot campaign, we collected a more extensive set of environmental parameters for each walk and observed correlations between the shoe sole dust microbial communities and the environmental conditions (Figure 6). Both the bacterial and fungal community compositions varied with temperature, PM_10_ and PM_2.5_ levels, and relative humidity, as well as whether the walking surface was wet or dry, visible dust was in the air, and snow was covering the ground (Table S2.b-h, S3.b-h). Some of these characteristics were highly positively correlated with each other (e.g. PM_10_ and PM_2.5_ and high visible dust, or relative humidity and wet surface), and others negatively (e.g., snow and temperature). These ordination plots also show how the conditions changed over the course of the pilot study as the year progressed from winter into spring. Environmental parameters showed significant correlation indicating potential co-influence on microbial community composition (Figure S8).

**Figure 6:**
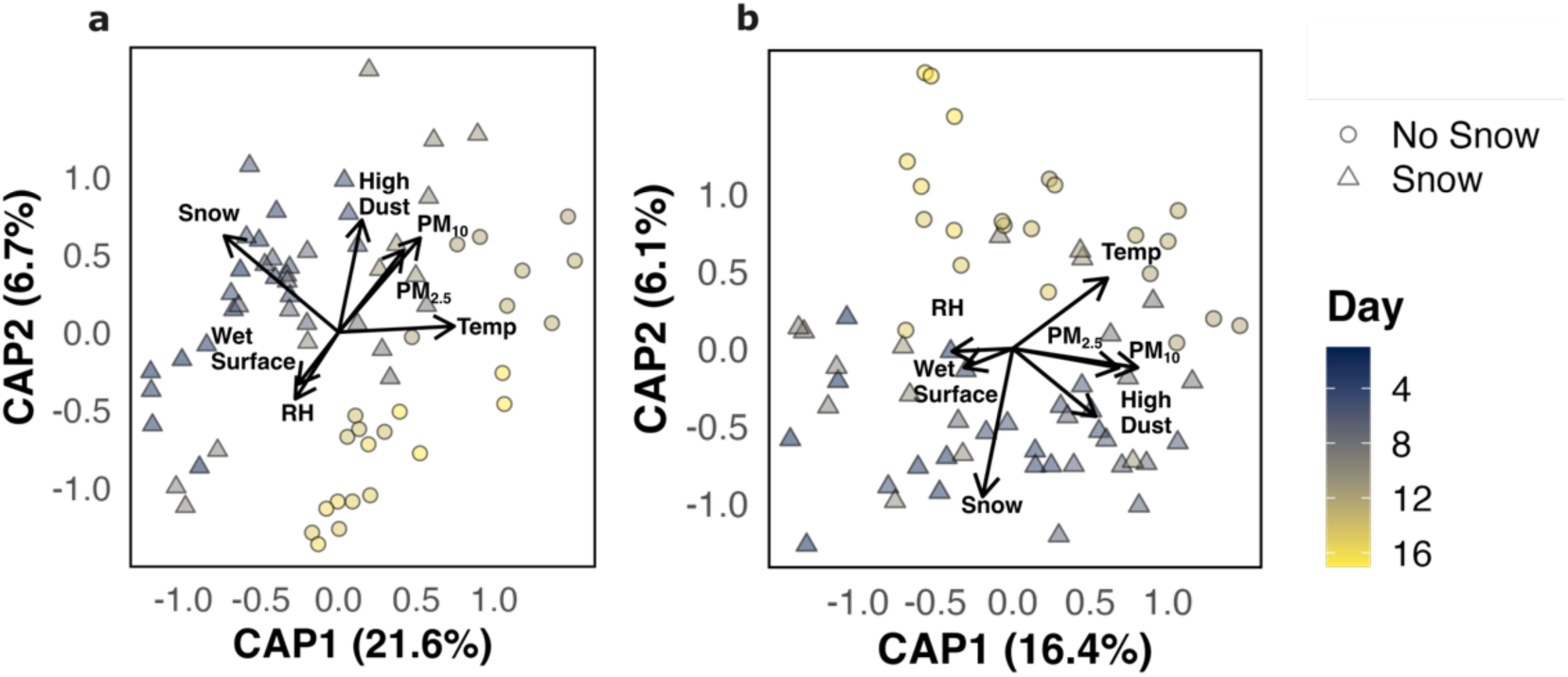
Constrained biplots showing the (a) bacterial and (b) fungal community composition at genus level in the pilot study explained by the environmental parameters. The circles represent samples collected when no snow was present on the ground, while triangles indicate samples collected when snow was present. Colors from dark blue to yellow illustrate the progression of sampling days from March to May.

### 4. Bacteria and fungi show differences in abundance and diversity between rural and urban environments

Both the bacterial and fungal communities from the shoe sole dust show clear differences along the urbanization gradient based on total biomass as measured by qPCR (Figure 7). Both total fungi and Gram-negative bacteria had the highest biomass in the rural-green samples. The total Gram-negative bacteria showed a clear stepwise pattern along the urbanization gradient from the high of the rural-green to the low of the urban-grey environment (p < 0.001for all comparisons), while total fungal biomass was similarly low in the urban-green and urban-grey environments (Figure 7). In contrast, the *Penicillium/Aspergillus* spp. group total biomass showed less clear differences across all three environments, with the highest abundance in urban-green and a statistically significant difference between the urban-grey and urban-green environments (p = 0.02). For total fungi, both the urban-grey and urban-green environments were significantly different from the rural-green environment (p < 0.001), but not from each other (Table S4).

**Figure 7:**
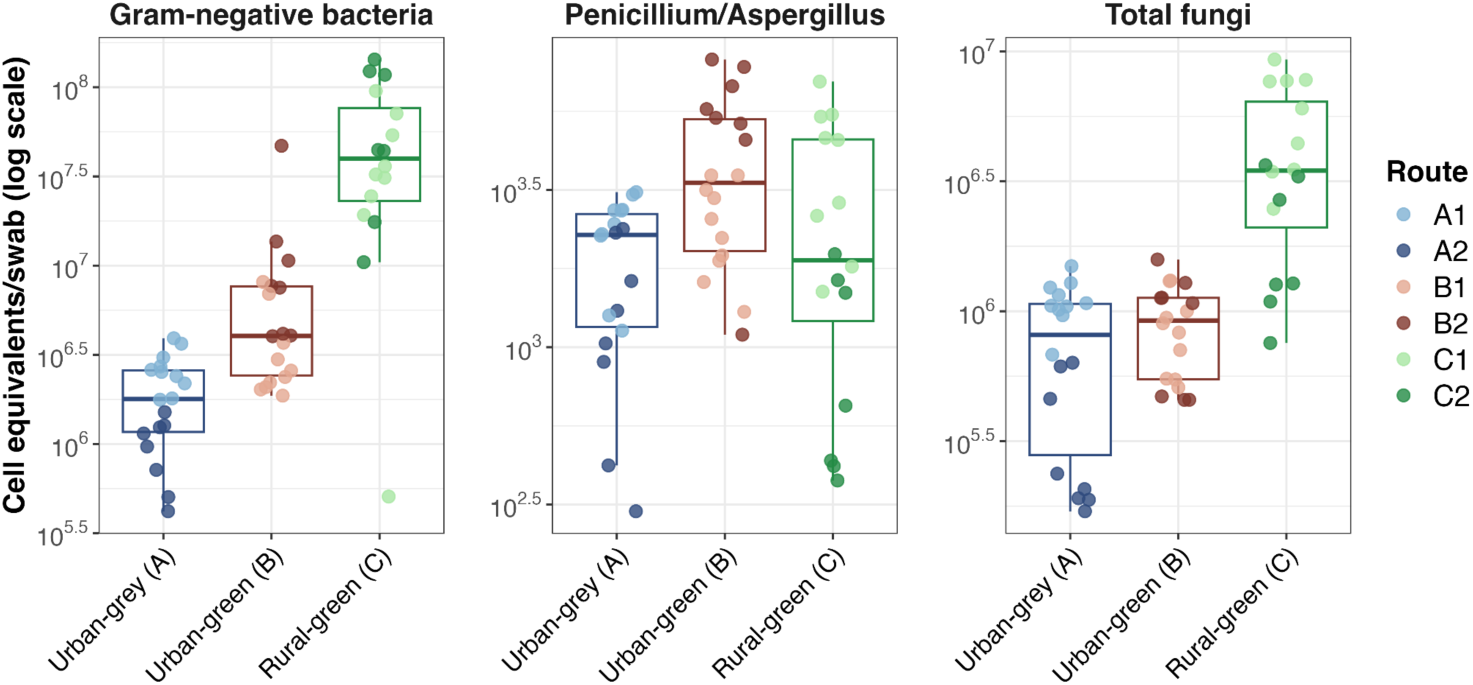
Biomass on shoe sole swabs as determined with qPCR targeting Gram-negative bacteria, Penicillium/Aspergillus group, and total fungi across the three environments (data presented with a log-scale). The boxplots are colored by the environment and the samples are colored by route.

The alpha diversity (both observed richness and Shannon diversity) of the fungal (ITS) and bacterial (16S) communities mirrored the patterns presented in the qPCR results. Alpha diversity was highest in the rural-green environment for both bacteria and fungi (Figure 8). Pairwise Wilcoxon tests revealed statistically significant differences in both observed richness and Shannon diversity between all environment pairs for the bacteria. Specifically, observed richness differed most significantly between the urban-grey and rural-green environments (p = 0.0004), followed by urban-grey versus urban-green (p = 0.015) and urban-green versus rural green (p = 0.025). Shannon diversity exhibited even stronger differences, with p < 0.001 for urban-grey versus both rural- and urban-green, and p < 0.0025 for rural-green versus urban-green. For fungi, Pairwise Wilcoxon tests confirmed significant differences in both observed richness and Shannon diversity, especially between the rural-green environment and both urban ones (p < 0.001 in all cases). On the other hand, there were no significant differences between the fungal alpha diversity in the urban-grey or urban-green environments.

**Figure 8:**
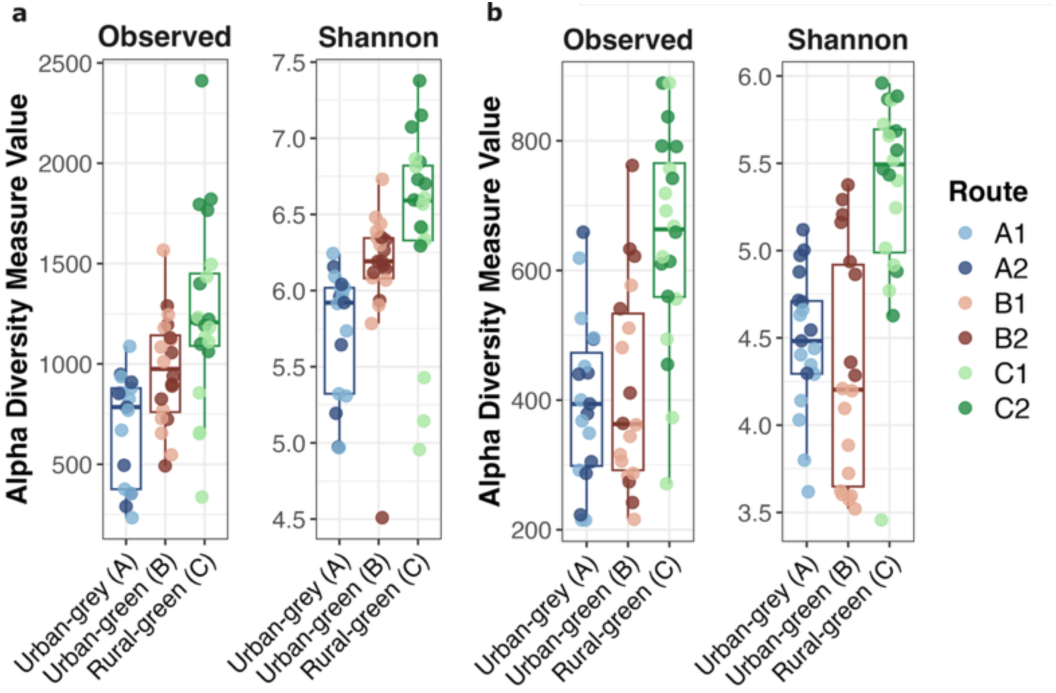
Alpha diversity (both observed ASV richness and Shannon dissimilarity) of (a) bacterial and (b) fungal communities showing significant differences across the three different environments. Here, the boxplots are colored by environment and the samples are colored by route. The y-axis shows the range of the diversity measure and does not start at zero to more clearly observe the differences. In all cases, the samples collected in the rural-green environment revealed the highest alpha diversity compared to others. Genus level plots show similar trends (Figure S9).

Environment also had a significant effect on the beta diversity of the bacterial (Table S5) and fungal (Table S6) communities determined from shoe sole dust (Figure 9). There was a clear difference between the rural-green and urban environments, with the urban routes clustering closely together and far away from the rural routes. More pronounced differences in the microbial communities were also apparent between the two rural-green routes (C1 and C2) than between any of the urban routes. Routes C1 and C2 and routes B1 and B2 were geographically similarly distant from each other (0.7-3.7km and 2.8-3.8km apart, respectively), while routes A1 and A2 were located much closer together (0.2-1 km apart) (Figure 1). For both the fungal and bacterial communities, the urban-grey routes were more similar compared to the urban-green routes. For fungi, the urban-green route B1 samples clustered more closely to the urban-grey samples than to the other urban green ones.

**Figure 9:**
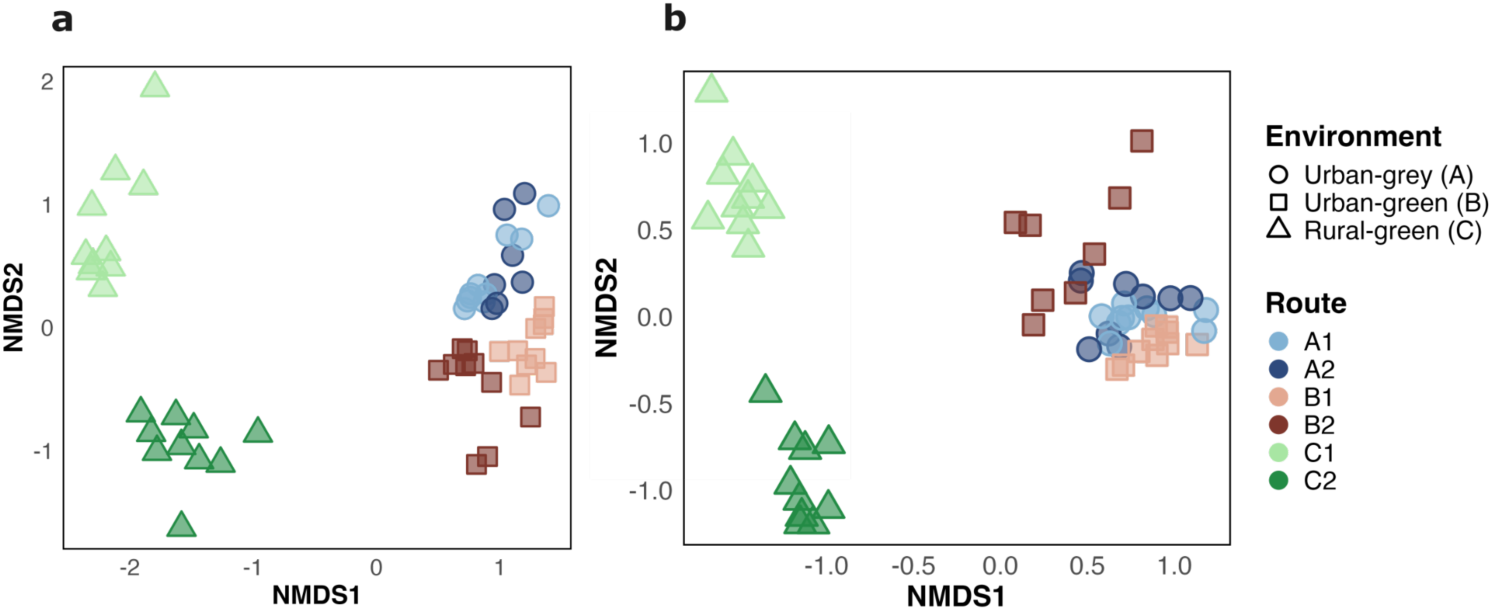
Beta diversity of (a) bacterial and (b) fungal communities across the three environments, visualized using Non-metric Multidimensional Scaling (NMDS) based on Bray-Curtis dissimilarity. The ordination plots reveal clear clustering patterns, indicating significant differences in microbial community composition between the environments for both bacterial (16S) and fungal (ITS) datasets. The NMDS stress values are 0.068 for the bacteria and 0.091 for fungi. Genus level plots are provided in Figure S10.

### 5. Bacterial diversity is more correlated with greenness closer to path than fungi

Both fungi and bacteria show clear increases in the number of taxa and Shannon diversity with increases in greenness along the routes as measured by NDVI (Figure 10; Table S7.a-g, S8.a-g). Figures S11-S17 show the plot of alpha diversity versus NDVI for buffers ranging from 10m to 10km. However, fungi and bacteria are differently dependent on the buffer size considered for calculating the NDVI. For bacteria, the correlations between greenness and the alpha diversity measures were strongest for the shortest buffer distances (10-50m), and the correlations decreased with increasing distance (up to 1000m). Conversely, the correlation for fungi were higher with a larger buffer area being used in the calculation of greenness (100-1000m) and lower correlations were seen for the smaller radii (10-50m).

**Figure 10:**
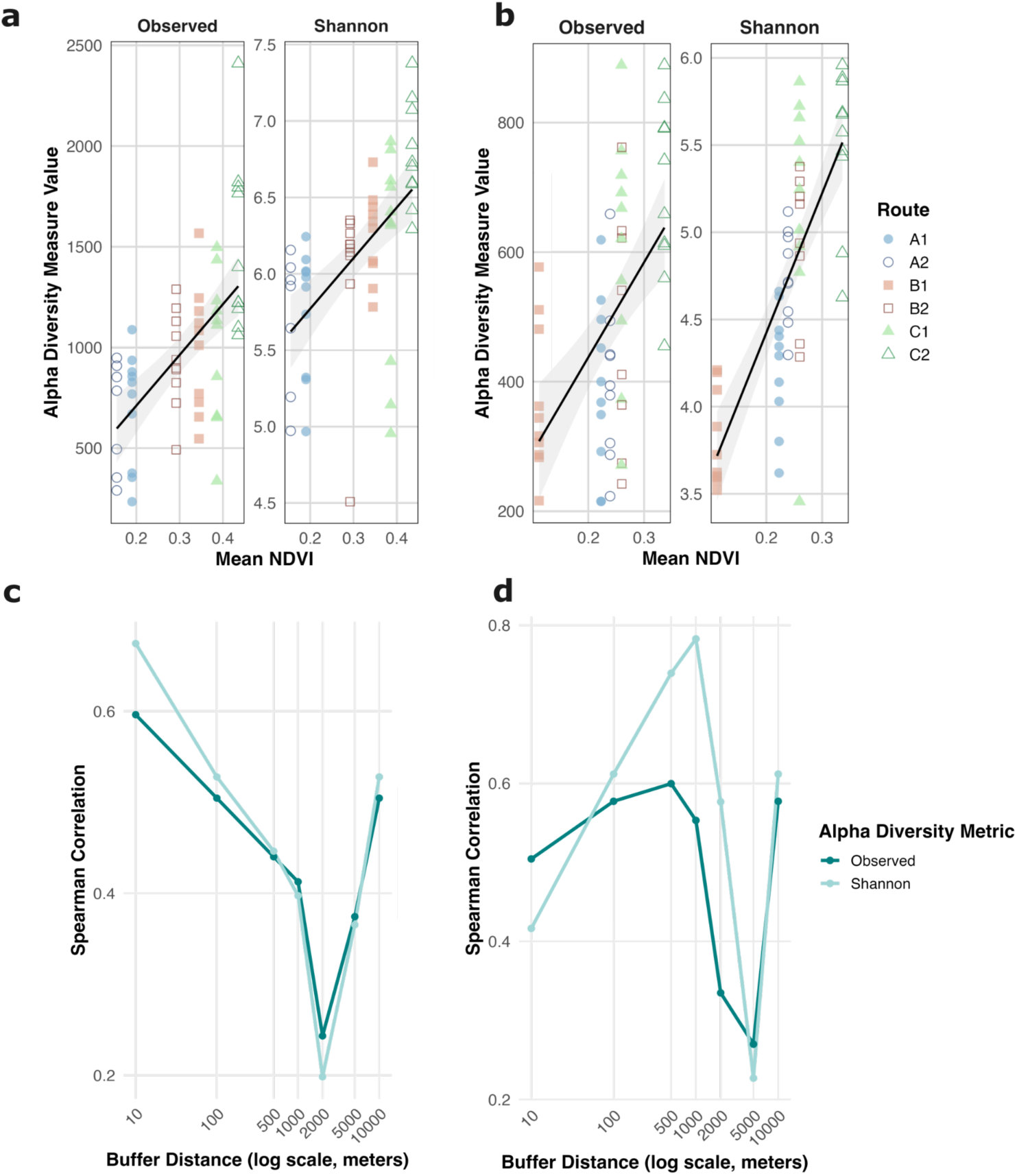
Correlation between alpha diversity of bacteria and fungi with mean NDVI: (a) alpha diversity (observed and Shannon) versus NDVI for the bacteria with a buffer of 10 m, (b) alpha diversity vs NDVI for the fungi at a buffer of 1 km. The bottom panel shows the plots of the correlations at each of the buffers (10-10,000m) for (c) bacteria and (d) fungi. Each point represents the Spearman correlation value between richness/diversity and NDVI for main study walks. Almost all correlations (25 out of 28) were statistically significant (Table S7.a-g, S8.a-g). As the same correlations were found for the buffer of 10m, 25m and 50m, as well as for 200m and 500m, we showed only 10m and 500m.

### 6. Specific microbes vary in abundance with urbanization level

For both bacterial and fungal communities, a high proportion of reads come from genera that are highly abundant in samples from both the urban-grey and urban-green environments (Figure 11). Very few genera were significantly different in relative abundance between the urban-grey (A) and urban-green (B) environments (Figure 12, Figure S18). Those genera that were differentially abundant between the urban (A and B) and rural (C) environments followed similar patterns for the fungi and bacteria. For both, there were larger log_2_F differences for the taxa that were enriched in the urban (A, B) environments than those that were enriched in the rural (C) environments. Additionally, similar taxa were differentially abundant in both urban environments compared to the rural environment, including fungi that were more abundant in the urban environments (e.g., *Knufia, Aureobasidium, Buckleyzyma, Filobasidium*) and less abundant (e.g., *Gyoerffyella*, *Solicoccoyma*, and *Chalara*), as well as bacteria that were more abundant (e.g., *Roseomonas*, *Microlunatas,* and *Acidophilium*) and less abundant (e.g., *Bryobacter* and *Jatrophihabitans*).

**Figure 11:**
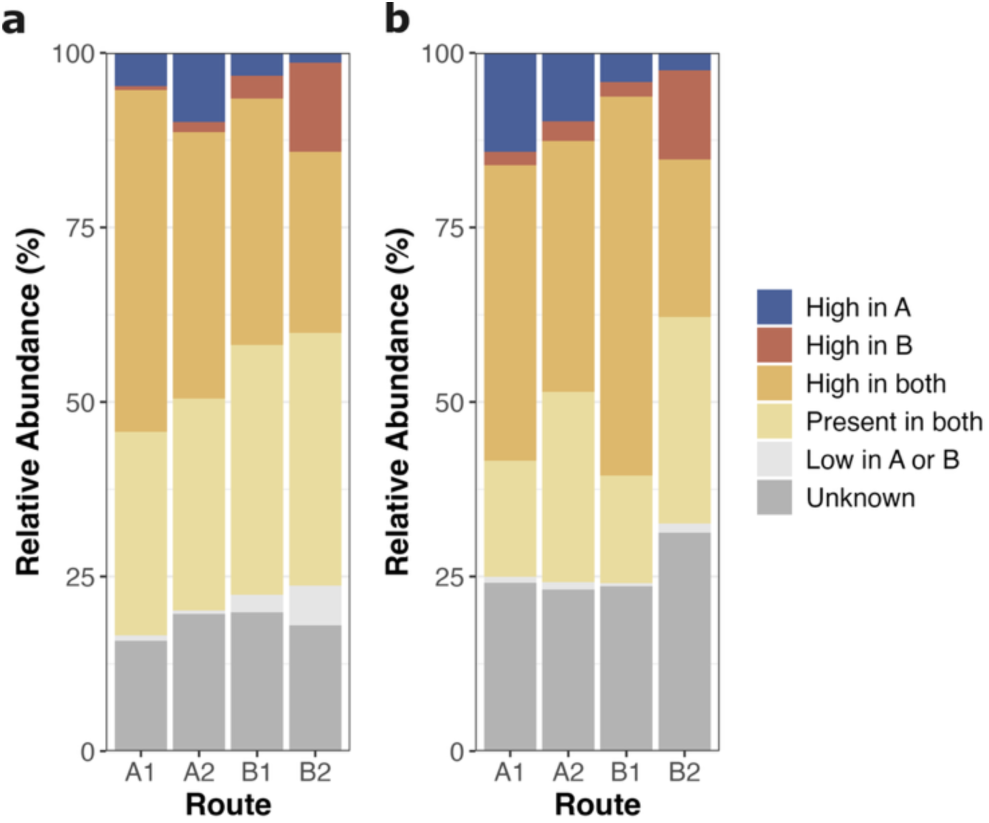
Comparison of genus-level relative abundances in shoe samples collected from the urban-grey and urban-green environments, separately for (a) bacterial and (b) fungal communities. Each bar chart is colored based on whether a particular genus was highly abundant (>1% relative abundance) only in samples from environment A (blue), only in samples from environment B (red), or whether it was highly abundant in both (orange). Yellow represents taxa that were present but not highly abundant in samples from both environments (<1% relative abundance), light grey marks taxa that were found at low levels in only one of the two environments. Dark grey (“Unknown”) represents reads belonging to ASVs that were not classified at the genus level.

**Figure 12:**
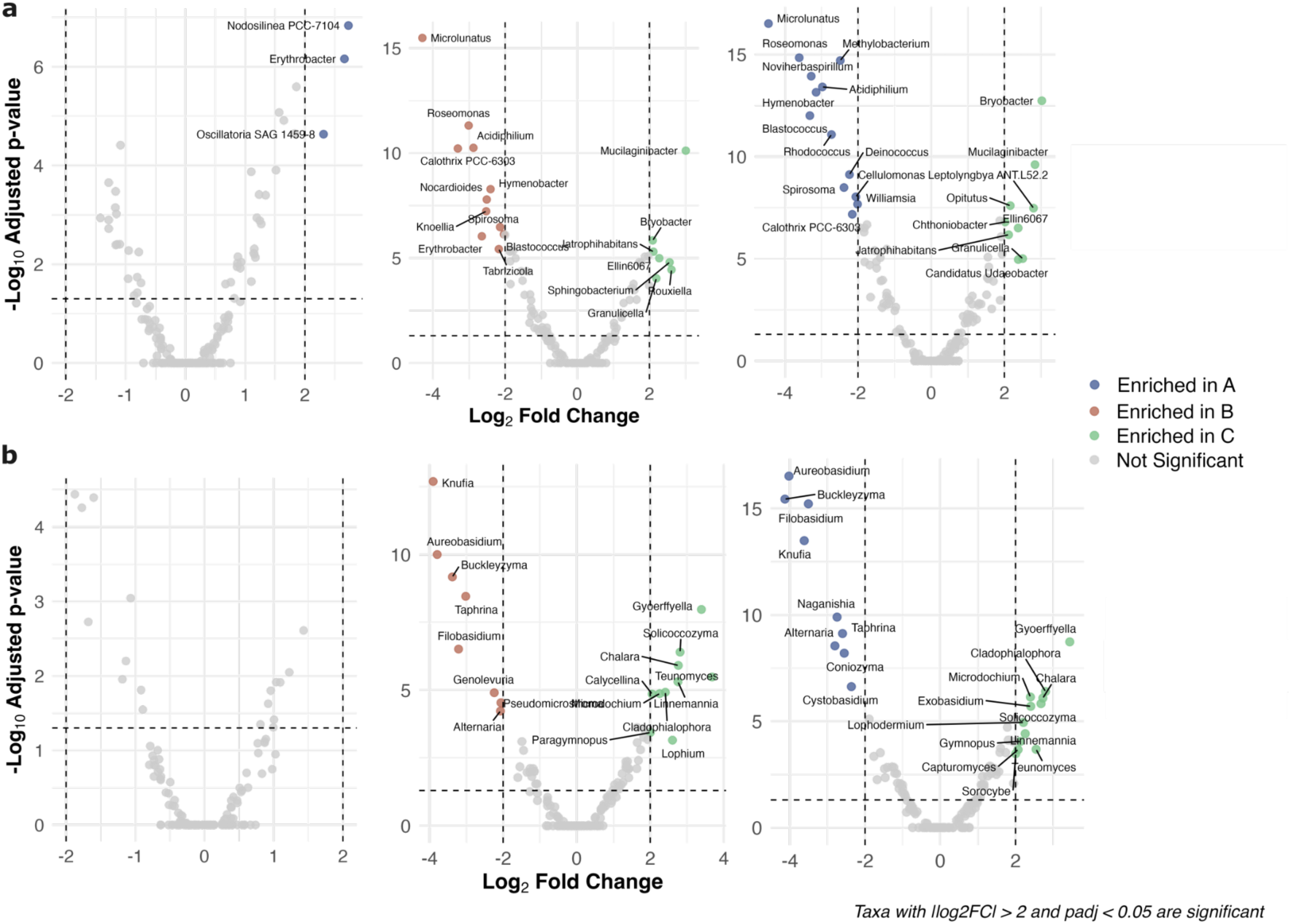
Volcano plots with statistically significant (a) bacterial and (b) fungal genera labeled. The negative log_10_ adjusted p-values are plotted versus the log_2_ fold changes. Left plots show urban-grey (A) versus urban-green (B), middle plots B versus rural-green (C), and right plots A versus C. Points are colored based on whether the taxa is enriched in environment A, B, or C.

## Discussion

### 1. Suitability and limitations of shoe sole dust for microbial exposure monitoring

In this study, we evaluated a simple and inexpensive approach for estimating microbial exposures during outdoor activities because of a lack of consideration of these exposures during epidemiological studies on asthma and allergies. Studies on the associations of indoor exposures and immunological development, asthma, and allergies commonly assess indoor microbial exposure through quantitative and qualitative microbial measurements from samples of indoor dust ^26,27,29,30^. Such approaches have facilitated breakthrough findings, including the beneficial effects of early childhood exposure to environmental microbes. However, comparable approaches that would allow assessment of microbial exposures during outdoor nature visits are not currently available. Assessing these exposures will require a simple approach akin to using settled dust for studies of indoor microbial exposure, therefore, we sought to validate whether shoe sole dust would reliably profile the microbial community of different outdoor activities. We are among the first to suggest and evaluate a simple and reliable approach for outdoor microbial exposure that could match the well-established protocols for those indoors.

We found that both fungal and bacterial diversity and biomass from the same walk (left and right shoe soles) were more similar than those from different walks on the same route or from walks on different routes. Our results show that while the left and right shoe soles do not return identical communities, they are similar to each other and reflect patterns consistent to the specific walk. We further demonstrate that the microbiota dissimilarity in shoe sole samples collected during walks on different routes is much higher than the dissimilarity between walks along the same route, which is higher than the dissimilarity for the same walk. We would not expect that the community on the left and right sole would be identical, as the paths they traversed are slightly different. Soil microbiomes vary greatly over small spatial scales, and even the left and right hands of the same individual vary in their microbiota reflecting similar differences in contacts ^60^. The observation that left and right samples from the same day have higher similarity compared to that of the same walk on a different day or a completely different walk demonstrates that these differences are small enough to not obscure other important biological differences. We also found that samples taken after walks in the morning and the afternoon of the same day (and the same route) were very similar, demonstrating that these microbial exposures were consistent over a single day. In addition to the large differences in our urban versus green environments and based on NDVI, we were also able to observe changes in the microbial communities based on environmental conditions during different walks on the same route.

We had consistently better DNA recoveries for the shoe sole swabs compared to the air samples. Only two air samples out of six were successfully sequenced for the fungal community and only one for the bacterial community. On the other hand, only three shoe sole dust samples (from after the walk) failed during sequencing for the bacterial community, and all passed for the fungal community, out of sixty samples. From the two walks that had both a shoe swab and air sample, we observed that the shoe sole swabs appeared to encompass most of the microbes in the air sample, as well as unique taxa. The taxa that were found in both the air and shoe sole samples suggest exchange between the ground and air microbial communities. The shoe sole sample’s broader detection may be due to the shoes’ contact with various microhabitats such as soil, leaf litter, and decaying organic matter, substrates that harbor a rich and diverse microbial pool not as readily dispersed into the air. Future studies, potentially in controlled chamber environments, would be necessary to comprehensively compare shoe sole dust to air samples during walking experiments on different surfaces, in order to ascertain the representativeness of shoe sole dust for airborne exposure to microbes. This will help establish when shoe sole dust versus air sampling is most appropriate. Assessing the benefits and drawbacks of a new sampling method is essential given the impact that this can have on the observed microbial community ^27,61–63^.

### 2. Nearby greenness impacts both fungal and bacterial community diversity captured on shoe soles, but in different ways

Both the fungal and bacterial communities of shoe sole dust showed higher diversity in the rural environment compared to the urban environments. NDVI, a proxy for nearby “greenness” (previously used by Fong et al., 2018 and Ju et al., 2024 to connect “greenness” quantitatively with health), explained the differences between our samples’ microbial communities better than either the environmental classification of “urban-green” vs “urban-grey” or the surface material of the path walked. Interestingly, the specific effect of greenness in the surrounding area was different for bacteria compared to fungi. Fungal diversity is more correlated with the greenness in a wider surrounding area (hundreds of meters), whereas bacterial diversity correlates better with greenness directly around the path (tens of meters). Route B1 exemplifies this trend: while it was an urban-green route based on the environment near to the route, its greenness was lower than the urban-grey routes for larger radii (100-1000 m), and samples collected along that route had the lowest fungal evenness. Considering different scales is essential for assessing the effect of greenness on human exposure to bacterial and fungal diversity; land use changes need to be coordinated at larger distances to increase the diversity of fungal versus bacterial exposures. These observations also indicate that a simple prediction of environmental microbial exposure based on land use or NDVI will likely not result in a reliable and sufficiently accurate exposure proxy, lending further support for approaches like shoe sole dust microbiota profiling in epidemiological studies.

The observation that bacterial and fungal communities respond differently to nearby land use suggests that different transport phenomena controlling bacterial and fungal communities in the air are at play. Fungal spores can disperse over longer distances through air ^65^, while bacterial cells are often more constrained by proximity to their source and environmental conditions ^66,67^. Similarly to our shoe sole dust results, a number of studies have reported that soil fungal richness was lowest at the highly urbanized site and highest at the lowly urbanized site^19–24^. Contrasting these results and ours, others have found that bacterial diversity declined significantly along the urban-to-rural gradient, while fungal diversity showed no significant variation ^25^. Reproducible methods like ours will help resolve whether these differences are related to study-specific effects (like sample collection) or reflective of different environments. Many studies report that urbanization causes urban areas to have microbial communities that are more similar to each other compared to rural areas ^20,24^, which we also observed in our study.

While Kuopio is a relatively green and low-pollution urban area compared to heavily urbanized regions, these differences still manifest clearly in the microbial communities. This suggests that even moderate urbanization can reduce microbial diversity and modify total biomass exposures, particularly for fungi. Based on these trends, we anticipate that more pronounced differences in microbial communities would be observed in larger, more densely urbanized environments.

### 3. Future directions and implications for human health

The patterns observed here are consistent with the mechanistic premise of the biodiversity hypothesis, which proposes that exposure to diverse environmental microbiota, particularly from natural ecosystems, can support healthy immune development ^8,68,69^. We clearly demonstrated higher microbial richness, diversity, and biomass in the rural-green environment compared to the urban environments. While total exposure to specific microbes, i.e., *Penicillium/Aspergillus*, was consistent across the three environments, their proportion of the total fungal DNA content dropped from the urban environments to rural ones. The sequencing data extends this pattern to other taxa, showing how the increase in microbial biomass in the rural environments is because of a gain in diversity rather than a pronounced increase in relative abundance of specific taxa. These results underscore how biodiversity loss and ecological imbalance in urban environments may favor less diverse communities, which in turn might influence human health, as proposed in the biodiversity hypothesis ^8,9,35,70^.

As the patterns connecting bacteria and fungi to nearby greenness diverge, it will be important for future public health interventions to understand how outdoor environmental exposure to bacteria and fungi are connected to health impacts. In a simplified example, if bacteria show stronger associations with health, then localized changes in greenness—such as tree planting and establishing small parks—may facilitate the desired health impacts. On the other hand, if fungal exposures were to show to be more relevant, then changes of greenness at a larger scale or actively entering rural environments will prove essential to achieve improvements in public health targets. Shoe sole dust microbial profiling may prove a useful tool in such future explorations.

While this study provides support of the mechanism behind the biodiversity hypothesis, the patterns observed here are correlational and did not include any health data. As this was a single study in one small city, more studies will be needed to test if these patterns persist in more and different urban environments or in different climatic zones. Future studies would be able to use this method of shoe sole sampling to associate health effects with the observed microbial diversity patterns. They will also be able to use this method to directly compare microbial exposures during different activities to assess the impact of lifestyle changes (e.g., spending more time in forest environments) on microbial exposures and health effects. For those looking to replicate this process in their own work, we have provided a detailed protocol at protocols.io ^39^. Our process is straightforward and would be simple for study participants to complete themselves following simple written or video instructions. Our work offers a simple approach for assessing outdoor exposures to complement the more common assessment of indoor microbiomes through settled dust samples that will allow for comprehensively accounting for both indoor and outdoor microbial exposures in epidemiological studies.

## Acknowledgements

Funding for this work comes from Dr. Hegarty’s start-up from the Case School of Engineering at CWRU and from the Finnish Institute for Health and Welfare. This research was supported by the Genomics Core Facility of the CWRU School of Medicine’s Genetics and Genome Sciences Department. We thank THL’s environmental microbiome lab for laboratory assistance in sample processing.

## References

(1) World Urbanization Prospects: The 2018 Revision; United Nations: New York, 2019.

(2) Adewuyi, F. A.; Knobel, P.; Gogna, P.; Dadvand, P. Health Effects of Green Prescription: A Systematic Review of Randomized Controlled Trials. Environ. Res. 2023, 236 (Pt 2), 116844. 10.1016/j.envres.2023.116844.

(3) Ccami-Bernal, F.; Soriano-Moreno, D. R.; Fernandez-Guzman, D.; Tuco, K. G.; Castro-Díaz, S. D.; Esparza-Varas, A. L.; Medina-Ramirez, S. A.; Caira-Chuquineyra, B.; Cortez-Soto, A. G.; Yovera-Aldana, M.; Rojas-Rueda, D. Green Space Exposure and Type 2 Diabetes Mellitus Incidence: A Systematic Review. Health Place 2023, 82, 103045. 10.1016/j.healthplace.2023.103045.

(4) Houlden, V.; Weich, S.; Porto de Albuquerque, J.; Jarvis, S.; Rees, K. The Relationship between Greenspace and the Mental Wellbeing of Adults: A Systematic Review. PloS One 2018, 13 (9), e0203000. 10.1371/journal.pone.0203000.

(5) Yang, B.-Y.; Zhao, T.; Hu, L.-X.; Browning, M. H. E. M.; Heinrich, J.; Dharmage, S. C.; Jalaludin, B.; Knibbs, L. D.; Liu, X.-X.; Luo, Y.-N.; James, P.; Li, S.; Huang, W.-Z.; Chen, G.; Zeng, X.-W.; Hu, L.-W.; Yu, Y.; Dong, G.-H. Greenspace and Human Health: An Umbrella Review. Innov. Camb. Mass 2021, 2 (4), 100164. 10.1016/j.xinn.2021.100164.

(6) Ruokolainen, L.; Parkkola, A.; Karkman, A.; Sinkko, H.; Peet, A.; Hämäläinen, A.-M.; von Hertzen, L.; Tillmann, V.; Koski, K.; Virtanen, S. M.; Niemelä, O.; Haahtela, T.; Knip, M. Contrasting Microbiotas between Finnish and Estonian Infants: Exposure to Acinetobacter May Contribute to the Allergy Gap. Allergy 2020, 75 (9), 2342–2351. 10.1111/all.14250.

(7) von Mutius, E.; Vercelli, D. Farm Living: Effects on Childhood Asthma and Allergy. Nat. Rev. Immunol. 2010, 10 (12), 861–868. 10.1038/nri2871.

(8) Haahtela, T. A Biodiversity Hypothesis. Allergy 2019, 74 (8), 1445–1456. 10.1111/all.13763.

(9) Hanski, I.; von Hertzen, L.; Fyhrquist, N.; Koskinen, K.; Torppa, K.; Laatikainen, T.; Karisola, P.; Auvinen, P.; Paulin, L.; Mäkelä, M. J.; Vartiainen, E.; Kosunen, T. U.; Alenius, H.; Haahtela, T. Environmental Biodiversity, Human Microbiota, and Allergy Are Interrelated. Proc. Natl. Acad. Sci. 2012, 109 (21), 8334–8339. 10.1073/pnas.1205624109.

(10) Ruokolainen, L.; Fyhrquist, N.; Haahtela, T. The Rich and the Poor: Environmental Biodiversity Protecting from Allergy. Curr. Opin. Allergy Clin. Immunol. 2016, 16 (5), 421–426. 10.1097/ACI.0000000000000304.

(11) Kirjavainen, P. V.; Karvonen, A. M.; Adams, R. I.; Täubel, M.; Roponen, M.; Tuoresmäki, P.; Loss, G.; Jayaprakash, B.; Depner, M.; Ege, M. J.; Renz, H.; Pfefferle, P. I.; Schaub, B.; Lauener, R.; Hyvärinen, A.; Knight, R.; Heederik, D. J. J.; von Mutius, E.; Pekkanen, J. Farm-like Indoor Microbiota in Non-Farm Homes Protects Children from Asthma Development. Nat. Med. 2019, 25 (7), 1089–1095. 10.1038/s41591-019-0469-4.

(12) Fuertes, E.; Markevych, I.; Bowatte, G.; Gruzieva, O.; Gehring, U.; Becker, A.; Berdel, D.; von Berg, A.; Bergström, A.; Brauer, M.; Brunekreef, B.; Brüske, I.; Carlsten, C.; Chan-Yeung, M.; Dharmage, S. C.; Hoffmann, B.; Klümper, C.; Koppelman, G. H.; Kozyrskyj, A.; Korek, M.; Kull, I.; Lodge, C.; Lowe, A.; MacIntyre, E.; Pershagen, G.; Standl, M.; Sugiri, D.; Wijga, A.; MACS; Heinrich, J. Residential Greenness Is Differentially Associated with Childhood Allergic Rhinitis and Aeroallergen Sensitization in Seven Birth Cohorts. Allergy 2016, 71 (10), 1461–1471. 10.1111/all.12915.

(13) Gernes, R.; Brokamp, C.; Rice, G. E.; Wright, J. M.; Kondo, M. C.; Michael, Y. L.; Donovan, G. H.; Gatziolis, D.; Bernstein, D.; LeMasters, G. K.; Lockey, J. E.; Khurana Hershey, G. K.; Ryan, P. H. Using High-Resolution Residential Greenspace Measures in an Urban Environment to Assess Risks of Allergy Outcomes in Children. Sci. Total Environ. 2019, 668, 760–767. 10.1016/j.scitotenv.2019.03.009.

(14) Lovasi, G. S.; O’Neil-Dunne, J. P. M.; Lu, J. W. T.; Sheehan, D.; Perzanowski, M. S.; Macfaden, S. W.; King, K. L.; Matte, T.; Miller, R. L.; Hoepner, L. A.; Perera, F. P.; Rundle, A. Urban Tree Canopy and Asthma, Wheeze, Rhinitis, and Allergic Sensitization to Tree Pollen in a New York City Birth Cohort. Environ. Health Perspect. 2013, 121 (4), 494–500. 10.1289/ehp.1205513.

(15) Lukkarinen, M.; Kirjavainen, P. V.; Backman, K.; Gonzales-Inca, C.; Hickman, B.; Kallio, S.; Karlsson, H.; Karlsson, L.; Keski-Nisula, L.; Korhonen, L. S.; Korpela, K.; Kuitunen, M.; Kukkonen, A. K.; Käyhkö, N.; Lagström, H.; Lukkarinen, H.; Peltola, V.; Pentti, J.; Salonen, A.; Savilahti, E.; Tuoresmäki, P.; Täubel, M.; Vahtera, J.; de Vos, W. M.; Pekkanen, J.; Karvonen, A. M. Early-Life Environment and the Risk of Eczema at 2 Years—Meta-Analyses of Six Finnish Birth Cohorts. Pediatr. Allergy Immunol. 2023, 34 (4), e13945. 10.1111/pai.13945.

(16) Parmes, E.; Pesce, G.; Sabel, C. E.; Baldacci, S.; Bono, R.; Brescianini, S.; D’Ippolito, C.; Hanke, W.; Horvat, M.; Liedes, H.; Maio, S.; Marchetti, P.; Marcon, A.; Medda, E.; Molinier, M.; Panunzi, S.; Pärkkä, J.; Polańska, K.; Prud’homme, J.; Ricci, P.; Snoj Tratnik, J.; Squillacioti, G.; Stazi, M. A.; Maesano, C. N.; Annesi-Maesano, I. Influence of Residential Land Cover on Childhood Allergic and Respiratory Symptoms and Diseases: Evidence from 9 European Cohorts. Environ. Res. 2020, 183, 108953. 10.1016/j.envres.2019.108953.

(17) Ruokolainen, L.; von Hertzen, L.; Fyhrquist, N.; Laatikainen, T.; Lehtomäki, J.; Auvinen, P.; Karvonen, A. M.; Hyvärinen, A.; Tillmann, V.; Niemelä, O.; Knip, M.; Haahtela, T.; Pekkanen, J.; Hanski, I. Green Areas around Homes Reduce Atopic Sensitization in Children. Allergy 2015, 70 (2), 195–202. 10.1111/all.12545.

(18) Tischer, C.; Gascon, M.; Fernández-Somoano, A.; Tardón, A.; Lertxundi Materola, A.; Ibarluzea, J.; Ferrero, A.; Estarlich, M.; Cirach, M.; Vrijheid, M.; Fuertes, E.; Dalmau-Bueno, A.; Nieuwenhuijsen, M. J.; Antó, J. M.; Sunyer, J.; Dadvand, P. Urban Green and Grey Space in Relation to Respiratory Health in Children. Eur. Respir. J. 2017, 49 (6), 1502112. 10.1183/13993003.02112-2015.

(19) Abrego, N.; Crosier, B.; Somervuo, P.; Ivanova, N.; Abrahamyan, A.; Abdi, A.; Hämäläinen, K.; Junninen, K.; Maunula, M.; Purhonen, J.; Ovaskainen, O. Fungal Communities Decline with Urbanization—More in Air than in Soil. ISME J. 2020, 14 (11), 2806–2815. 10.1038/s41396-020-0732-1.

(20) Barberán, A.; Ladau, J.; Leff, J. W.; Pollard, K. S.; Menninger, H. L.; Dunn, R. R.; Fierer, N. Continental-Scale Distributions of Dust-Associated Bacteria and Fungi. Proc. Natl. Acad. Sci. 2015, 112 (18), 5756–5761. 10.1073/pnas.1420815112.

(21) Chen, Y.; Martinez, A.; Cleavenger, S.; Rudolph, J.; Barberán, A. Changes in Soil Microbial Communities across an Urbanization Gradient: A Local-Scale Temporal Study in the Arid Southwestern USA. Microorganisms 2021, 9 (7), 1470. 10.3390/microorganisms9071470.

(22) Epp Schmidt, D. J.; Pouyat, R.; Szlavecz, K.; Setälä, H.; Kotze, D. J.; Yesilonis, I.; Cilliers, S.; Hornung, E.; Dombos, M.; Yarwood, S. A. Urbanization Erodes Ectomycorrhizal Fungal Diversity and May Cause Microbial Communities to Converge. Nat. Ecol. Evol. 2017, 1 (5), 123. 10.1038/s41559-017-0123.

(23) Flies, E. J.; Clarke, L. J.; Brook, B. W.; Jones, P. Urbanisation Reduces the Abundance and Diversity of Airborne Microbes - but What Does That Mean for Our Health? A Systematic Review. Sci. Total Environ. 2020, 738, 140337. 10.1016/j.scitotenv.2020.140337.

(24) Zheng, B.; Hui, N.; Jumpponen, A.; Lu, C.; Pouyat, R.; Szlavecz, K.; Wardle, D. A.; Yesilonis, I.; Setälä, H.; Kotze, D. J. Urbanization Leads to Asynchronous Homogenization of Soil Microbial Communities across Biomes. Environ. Sci. Ecotechnology 2025, 25, 100547. 10.1016/j.ese.2025.100547.

(25) Tan, X.; Kan, L.; Su, Z.; Liu, X.; Zhang, L. The Composition and Diversity of Soil Bacterial and Fungal Communities Along an Urban-To-Rural Gradient in South China. Forests 2019, 10 (9), 797. 10.3390/f10090797.

(26) Adams, R. I.; Tian, Y.; Taylor, J. W.; Bruns, T. D.; Hyvärinen, A.; Täubel, M. Passive Dust Collectors for Assessing Airborne Microbial Material. Microbiome 2015, 3, 46. 10.1186/s40168-015-0112-7.

(27) Frankel, M.; Timm, M.; Hansen, E. W.; Madsen, A. M. Comparison of Sampling Methods for the Assessment of Indoor Microbial Exposure. Indoor Air 2012, 22 (5), 405–414. 10.1111/j.1600-0668.2012.00770.x.

(28) Institute of Medicine (US) Committee on Damp Indoor Spaces and Health. Damp Indoor Spaces and Health; National Academies Press (US): Washington (DC), 2004.

(29) Leppänen, H. K.; Täubel, M.; Jayaprakash, B.; Vepsäläinen, A.; Pasanen, P.; Hyvärinen, A. Quantitative Assessment of Microbes from Samples of Indoor Air and Dust. J. Expo. Sci. Environ. Epidemiol. 2018, 28 (3), 231–241. 10.1038/jes.2017.24.

(30) Noss, I.; Wouters, I. M.; Visser, M.; Heederik, D. J. J.; Thorne, P. S.; Brunekreef, B.; Doekes, G. Evaluation of a Low-Cost Electrostatic Dust Fall Collector for Indoor Air Endotoxin Exposure Assessment. Appl. Environ. Microbiol. 2008, 74 (18), 5621–5627. 10.1128/AEM.00619-08.

(31) Turunen, A. W.; Halonen, J.; Korpela, K.; Ojala, A.; Pasanen, T.; Siponen, T.; Tiittanen, P.; Tyrväinen, L.; Yli-Tuomi, T.; Lanki, T. Cross-Sectional Associations of Different Types of Nature Exposure with Psychotropic, Antihypertensive and Asthma Medication. Occup. Environ. Med. 2023, 80 (2), 111–118. 10.1136/oemed-2022-108491.

(32) Bardhan, M.; Zhang, K.; Browning, M. H. E. M.; Dong, J.; Liu, T.; Bailey, C.; McAnirlin, O.; Hanley, J.; Minson, C. T.; Mutel, R. L.; Ranganathan, S.; Reuben, A. Time in Nature Is Associated with Higher Levels of Positive Mood: Evidence from the 2023 NatureDose^TM^ Student Survey. J. Environ. Psychol. 2023, 90, 102083. 10.1016/j.jenvp.2023.102083.

(33) MacKerron, G.; Mourato, S. Happiness Is Greater in Natural Environments. Glob. Environ. Change 2013, 23 (5), 992–1000. 10.1016/j.gloenvcha.2013.03.010.

(34) Winnicki, M. H.; Dunn, R. R.; Winther-Jensen, M.; Jess, T.; Allin, K. H.; Bruun, H. H. Does Childhood Exposure to Biodiverse Greenspace Reduce the Risk of Developing Asthma? Sci. Total Environ. 2022, 850, 157853. 10.1016/j.scitotenv.2022.157853.

(35) Potter, J. D.; Brooks, C.; Donovan, G.; Cunningham, C.; Douwes, J. A Perspective on Green, Blue, and Grey Spaces, Biodiversity, Microbiota, and Human Health. Sci. Total Environ. 2023, 892, 164772. 10.1016/j.scitotenv.2023.164772.

(36) Lax, S.; Hampton-Marcell, J. T.; Gibbons, S. M.; Colares, G. B.; Smith, D.; Eisen, J. A.; Gilbert, J. A. Forensic Analysis of the Microbiome of Phones and Shoes. Microbiome 2015, 3 (1), 21. 10.1186/s40168-015-0082-9.

(37) Coil, D. A.; Neches, R. Y.; Lang, J. M.; Jospin, G.; Brown, W. E.; Cavalier, D.; Hampton-Marcell, J.; Gilbert, J. A.; Eisen, J. A. Bacterial Communities Associated with Cell Phones and Shoes. PeerJ 2020, 8, e9235. 10.7717/peerj.9235.

(38) Salonen, R.; Siponen, T.; Taimisto, P.; Pärjälä, E. Katupölylle altistuminen Kuopion keskustassa keväällä 2019. 2019.

(39) Täubel, M.; Ferdous, S. M.; Hegarty, B. Shoe Sole Dust Sampling Protocol. 2025.

(40) Haugland, R. A.; Siefring, S. C.; Wymer, L. J.; Brenner, K. P.; Dufour, A. P. Comparison of Enterococcus Measurements in Freshwater at Two Recreational Beaches by Quantitative Polymerase Chain Reaction and Membrane Filter Culture Analysis. Water Res. 2005, 39 (4), 559–568. 10.1016/j.watres.2004.11.011.

(41) Haugland, R. A.; Brinkman, N.; Vesper, S. J. Evaluation of Rapid DNA Extraction Methods for the Quantitative Detection of Fungi Using Real-Time PCR Analysis. J. Microbiol. Methods 2002, 50 (3), 319–323. 10.1016/s0167-7012(02)00037-4.

(42) Hyytiäinen, H. K.; Jayaprakash, B.; Kirjavainen, P. V.; Saari, S. E.; Holopainen, R.; Keskinen, J.; Hämeri, K.; Hyvärinen, A.; Boor, B. E.; Täubel, M. Crawling-Induced Floor Dust Resuspension Affects the Microbiota of the Infant Breathing Zone. Microbiome 2018, 6 (1), 25. 10.1186/s40168-018-0405-8.

(43) Kärkkäinen, P. M.; Valkonen, M.; Hyvärinen, A.; Nevalainen, A.; Rintala, H. Determination of Bacterial Load in House Dust Using qPCR, Chemical Markers and Culture. J. Environ. Monit. JEM 2010, 12 (3), 759–768. 10.1039/b917937b.

(44) Haugland, R. A.; Varma, M.; Wymer, L. J.; Vesper, S. J. Quantitative PCR Analysis of Selected Aspergillus, Penicillium and Paecilomyces Species. Syst. Appl. Microbiol. 2004, 27 (2), 198–210. 10.1078/072320204322881826.

(45) Haugland, R.; Vesper, S. J. Method of Identifying and Quantifying Specific Fungi and Bacteria. WO2001096612A2, December 20, 2001. https://patents.google.com/patent/WO2001096612A2/en (accessed 2025-11-25).

(46) Caporaso, J. G.; Lauber, C. L.; Walters, W. A.; Berg-Lyons, D.; Lozupone, C. A.; Turnbaugh, P. J.; Fierer, N.; Knight, R. Global Patterns of 16S rRNA Diversity at a Depth of Millions of Sequences per Sample. Proc. Natl. Acad. Sci. U. S. A. 2011, 108 Suppl 1 (Suppl 1), 4516–4522. 10.1073/pnas.1000080107.

(47) Smith, D. P.; Peay, K. G. Sequence Depth, Not PCR Replication, Improves Ecological Inference from next Generation DNA Sequencing. PloS One 2014, 9 (2), e90234. 10.1371/journal.pone.0090234.

(48) Dockx, Y.; Täubel, M.; Bijnens, E. M.; Witters, K.; Valkonen, M.; Jayaprakash, B.; Hogervorst, J.; Nawrot, T. S.; Casas, L. Residential Green Space Can Shape the Indoor Microbial Environment. Environ. Res. 2021, 201, 111543. 10.1016/j.envres.2021.111543.

(49) Klindworth, A.; Pruesse, E.; Schweer, T.; Peplies, J.; Quast, C.; Horn, M.; Glöckner, F. O. Evaluation of General 16S Ribosomal RNA Gene PCR Primers for Classical and Next-Generation Sequencing-Based Diversity Studies. Nucleic Acids Res. 2013, 41 (1), e1. 10.1093/nar/gks808.

(50) White, T.; Bruns, T.; Taylor, J. 38 - Amplification And Direct Sequencing of Fungal Ribosomal RNA Genes For Phylogenetics. In PCR Protocols: A Guide to Methods and Applications; Academic Press: USA, 1990; pp 315–322.

(51) Leslie, A.; Chowdhury, M. R.; Täubel, M.; Hegarty, B. Indoor Potted Plants Have Little Effect on Office Dust Fungal Communities. Indoor Environ. 2025, 2 (2), 100092. 10.1016/j.indenv.2025.100092.

(52) Callahan, B. J.; McMurdie, P. J.; Holmes, S. P. Exact Sequence Variants Should Replace Operational Taxonomic Units in Marker-Gene Data Analysis. ISME J. 2017, 11 (12), 2639–2643. 10.1038/ismej.2017.119.

(53) Quast, C.; Pruesse, E.; Yilmaz, P.; Gerken, J.; Schweer, T.; Yarza, P.; Peplies, J.; Glöckner, F. O. The SILVA Ribosomal RNA Gene Database Project: Improved Data Processing and Web-Based Tools. Nucleic Acids Res. 2013, 41 (Database issue), D590–596. 10.1093/nar/gks1219.

(54) Abarenkov, K.; Nilsson, R. H.; Larsson, K.-H.; Taylor, A. F. S.; May, T. W.; Frøslev, T. G.; Pawlowska, J.; Lindahl, B.; Põldmaa, K.; Truong, C.; Vu, D.; Hosoya, T.; Niskanen, T.; Piirmann, T.; Ivanov, F.; Zirk, A.; Peterson, M.; Cheeke, T. E.; Ishigami, Y.; Jansson, A. T.; Jeppesen, T. S.; Kristiansson, E.; Mikryukov, V.; Miller, J. T.; Oono, R.; Ossandon, F. J.; Paupério, J.; Saar, I.; Schigel, D.; Suija, A.; Tedersoo, L.; Kõljalg, U. The UNITE Database for Molecular Identification and Taxonomic Communication of Fungi and Other Eukaryotes: Sequences, Taxa and Classifications Reconsidered. Nucleic Acids Res. 2024, 52 (D1), D791–D797. 10.1093/nar/gkad1039.

(55) Poulsen, C. S.; Kaas, R. S.; Aarestrup, F. M.; Pamp, S. J. Standard Sample Storage Conditions Have an Impact on Inferred Microbiome Composition and Antimicrobial Resistance Patterns. Microbiol. Spectr. 2021, 9 (2), e01387–21. 10.1128/Spectrum.01387-21.

(56) Copernicus Sentinel Data. European Union/ESA/Copernicus, 2020. https://browser.dataspace.copernicus.eu/ (accessed 2025-06-26).

(57) Rhew, I. C.; Vander Stoep, A.; Kearney, A.; Smith, N. L.; Dunbar, M. D. Validation of the Normalized Difference Vegetation Index as a Measure of Neighborhood Greenness. Ann. Epidemiol. 2011, 21 (12), 946–952. 10.1016/j.annepidem.2011.09.001.

(58) McMurdie, P. J.; Holmes, S. Phyloseq: An R Package for Reproducible Interactive Analysis and Graphics of Microbiome Census Data. PloS One 2013, 8 (4), e61217. 10.1371/journal.pone.0061217.

(59) Lin, H.; Peddada, S. D. Multigroup Analysis of Compositions of Microbiomes with Covariate Adjustments and Repeated Measures. Nat. Methods 2024, 21 (1), 83–91. 10.1038/s41592-023-02092-7.

(60) Fierer, N.; Hamady, M.; Lauber, C. L.; Knight, R. The Influence of Sex, Handedness, and Washing on the Diversity of Hand Surface Bacteria. Proc. Natl. Acad. Sci. U. S. A. 2008, 105 (46), 17994–17999. 10.1073/pnas.0807920105.

(61) Cox, J.; Indugula, R.; Vesper, S.; Zhu, Z.; Jandarov, R.; Reponen, T. Comparison of Indoor Air Sampling and Dust Collection Methods for Fungal Exposure Assessment Using Quantitative PCR. Environ. Sci. Process. Impacts 2017, 19 (10), 1312–1319. 10.1039/c7em00257b.

(62) Hoisington, A. J.; Maestre, J. P.; King, M. D.; Siegel, J. A.; Kinney, K. A. Impact of Sampler Selection on the Characterization of the Indoor Microbiome via High-Throughput Sequencing. Build. Environ. 2014, 80, 274–282. 10.1016/j.buildenv.2014.04.021.

(63) Knudsen, B. E.; Bergmark, L.; Munk, P.; Lukjancenko, O.; Priemé, A.; Aarestrup, F. M.; Pamp, S. J. Impact of Sample Type and DNA Isolation Procedure on Genomic Inference of Microbiome Composition. mSystems 2016, 1 (5), 10.1128/msystems.00095-16. 10.1128/msystems.00095-16.

(64) Ju, Y.; Dronova, I.; Ma, Q.; Lin, J.; Moran, M. R.; Gouveia, N.; Hu, H.; Yin, H.; Shang, H. Assessing Normalized Difference Vegetation Index as a Proxy of Urban Greenspace Exposure. Urban For. Urban Green. 2024, 99, 128454. 10.1016/j.ufug.2024.128454.

(65) Golan, J. J.; Pringle, A. Long-Distance Dispersal of Fungi. Microbiol. Spectr. 2017, 5 (4), 10.1128/microbiolspec.funk-0047–2016. 10.1128/microbiolspec.funk-0047-2016.

(66) Tong, X.; Leung, M. H. Y.; Wilkins, D.; Lee, P. K. H. City-Scale Distribution and Dispersal Routes of Mycobiome in Residences. Microbiome 2017, 5 (1), 131. 10.1186/s40168-017-0346-7.

(67) Walters, K. E.; Capocchi, J. K.; Albright, M. B. N.; Hao, Z.; Brodie, E. L.; Martiny, J. B. H. Routes and Rates of Bacterial Dispersal Impact Surface Soil Microbiome Composition and Functioning. ISME J. 2022, 16 (10), 2295–2304. 10.1038/s41396-022-01269-w.

(68) Haahtela, T.; Holgate, S.; Pawankar, R.; Akdis, C. A.; Benjaponpitak, S.; Caraballo, L.; Demain, J.; Portnoy, J.; von Hertzen, L.; WAO Special Committee on Climate Change and Biodiversity. The Biodiversity Hypothesis and Allergic Disease: World Allergy Organization Position Statement. World Allergy Organ. J. 2013, 6 (1), 3. 10.1186/1939-4551-6-3.

(69) Marselle, M. R.; Hartig, T.; Cox, D. T. C.; de Bell, S.; Knapp, S.; Lindley, S.; Triguero-Mas, M.; Böhning-Gaese, K.; Braubach, M.; Cook, P. A.; de Vries, S.; Heintz-Buschart, A.; Hofmann, M.; Irvine, K. N.; Kabisch, N.; Kolek, F.; Kraemer, R.; Markevych, I.; Martens, D.; Müller, R.; Nieuwenhuijsen, M.; Potts, J. M.; Stadler, J.; Walton, S.; Warber, S. L.; Bonn, A. Pathways Linking Biodiversity to Human Health: A Conceptual Framework. Environ. Int. 2021, 150, 106420. 10.1016/j.envint.2021.106420.

(70) Haahtela, T.; Alenius, H.; Lehtimäki, J.; Sinkkonen, A.; Fyhrquist, N.; Hyöty, H.; Ruokolainen, L.; Mäkelä, M. J. Immunological Resilience and Biodiversity for Prevention of Allergic Diseases and Asthma. Allergy 2021, 76 (12), 3613–3626. 10.1111/all.14895.

